# Combined statin–probiotic treatment mitigates atherogenic dyslipidemia fueling NAFLD in high-cholesterol-diet fed rats

**DOI:** 10.64898/2026.10.15.759314

**Authors:** Abinayaa Rajkumar, Priyadharshini Elangovan, Gokulprasanth Panchalingam, Devasimman Perumal, Vijayakrishnan Sevel Rangaswamy, Kalaiselvi Periandavan

## Abstract

**Background:** Chronic hypercholesterolemia accelerates metabolic dysregulation, systemic oxidative stress, and endothelial dysfunction, driving the initiation and progression of atherogenesis. Although statins like Atorvastatin (AS) represent the clinical cornerstone of lipid-lowering therapy, long-term high-dose use is often constrained by adverse side effects. Probiotic interventions targeting the gut–heart axis, such as *Bifidobacterium longum* (B), offer a promising therapeutic strategy to enhance lipid homeostasis and reduce vascular inflammation.

**Methods:** Male rats were fed a High-Cholesterol Diet (HCD) for 120 days to establish hypercholesterolemia, dyslipidemia, and systemic vascular pathology. Experimental groups were administered AS monotherapy, B. longum monotherapy, or their dual combination (AS + B) daily from day 90 to 120. Redox dynamics (GSH, MDA, GSH/MDA ratio), functional HDL-associated proteins (PON1, APOA1), circulating oxidized LDL (oxLDL), inflammatory cytokines (CRP, TNF-α, IL-6, IL-8), and all adhesion molecules (ICAM-1, VCAM-1) were quantified, alongside histopathological evaluation of hepatic and aortic tissue microarchitecture.

**Results:** High-cholesterol diet feeding induced severe dyslipidemia, vascular remodelling, lipid peroxidation, and endothelial inflammation in rats. Monotherapy with Atorvastatin (AS) or *Bifidobacterium longum* (B) significantly reversed metabolic impairment (p<0.001), while dual AS + B therapy achieved maximum efficacy (p<0.01 vs. AS alone) in restoring hepatic and aortic microarchitecture. Mechanistically, combination treatment blunted oxidative stress by minimizing malondialdehyde and upregulating Paraoxonase 1 and Apolipoprotein A1 (p<0.01), which markedly suppressed low-density lipoprotein oxidation (p<0.001). Furthermore, dual therapy yielded the maximum downregulation of pro-inflammatory cytokines (C-reactive protein, tumor necrosis factor-α, interleukin-6, interleukin-8) and adhesion molecules (ICAM1,VCAM1); p<0.001), demonstrating potent anti-inflammatory effect against diet-induced atherogenesis.

**Conclusion:** Co-administration of Atorvastatin and *Bifidobacterium longum* demonstrates multi-targeted therapeutic synergy by effectively restoring HDL functional capacity, attenuating vascular endothelial inflammation, and mitigating hypercholesterolemia simultaneously, this dual pharmacological–probiotic strategy provides a robust, novel intervention against diet-induced atherogenesis.

## INTRODUCTION

Cardiovascular diseases (CVDs) represent a major global health burden accounting for 17.9 million deaths annually and remains the leading cause of morbidity and mortality worldwide. Among CVD manifestations, atherosclerosis, a chronic inflammatory lifestyle disease characterized by the progressive accumulation of lipids and fibrous elements in large arteries which constitutes 50% of all deaths in westernized societies (Pahwa and Jialal, 2023; Rajkumar *et al*., 2025). Because atherosclerosis follows a silent progression pattern, it typically remains clinically asymptomatic until reaching advanced stages (Luca *et al*., 2023), frequently resulting in acute vascular events, poor prognoses, and constrained treatment outcomes.

Diet-induced metabolic disturbances, particularly those arising from high-fat and high-cholesterol consumption, play a central role in driving atherogenesis (Lordan *et al*., 2019). Chronic high-fat intake severely disrupts systemic lipid homeostasis, and these alterations accelerate vascular lesion formation through subendothelial lipid deposition, oxidative modification, and inflammatory activation (Rajkumar *et al*., 2025). Concurrently, hypercholesterolemia drives systemic oxidative stress characterized by excessive reactive oxygen species (ROS) generation and compromised antioxidant defenses which promotes lipid peroxidation, endothelial injury, and vascular remodeling, initiating atherogenic dyslipidemia, a pro-inflammatory state and a pro-thrombotic state thereby amplifying overall cardiovascular risk (Lasker *et al*., 2019). Pharmacological interventions, particularly statin therapy, remain the cornerstone of dyslipidemia and clinical atherosclerosis management (Grundy et al., 2019). Atorvastatin, a potent 3-hydroxy-3-methylglutaryl-coenzyme A (HMG-CoA) reductase inhibitor, effectively lowers LDL-C levels and improves overall lipid profiles. Beyond its primary lipid-lowering capacity, Atorvastatin exerts well-recognized pleiotropic benefits, including anti-inflammatory, plaque-stabilizing, and antioxidant actions that contribute to vascular preservation. However, concerns regarding long-term, high-dose statin administration including potential statin intolerance, myopathies, and hepatic enzyme elevations (Ray, 2024) have prompted intense interest in alternative or complementary therapeutic strategies to optimize clinical outcomes while minimizing dosage-related adverse effects.

In recent years, the “gut-heart axis” has emerged as a fundamental physiological network connecting dietary components, intestinal microbial ecology, and cardiovascular homeostasis. The gut microbiota functions as an essential bridge between diet and host physiology by metabolizing ingested substrates into diverse, bioactive metabolites (Tang and coworkers, 2017). Key gut-derived metabolites, such as short-chain fatty acids (SCFAs), conjugated fatty acids, secondary bile acids, and trimethylamine N-oxide (TMAO), act as distant signaling molecules that modulate systemic inflammation, vascular tone, and lipid absorption (Pieczynska et al 2020). Gut microbiota derived metabolites interact with cardiovascular phenotypes in various molecular pathways and host receptors. (Abdulrahim *et al*., 2025) Consequently, gut microbiota-targeted interventions have gained significant attraction as novel metabolic therapeutics. Probiotics are defined by the World Health Organization (WHO) as “live microorganisms which when administered in adequate amounts confer a health benefit on the host” (Sugahara et al., 2015). Probiotics exert systemic protective effects primarily by modulating host immunological responses, regulating inflammatory cascades through SCFA secretion, competing for mucosal adhesion, pathogenic suppression and preserve intestinal barrier integrity (Suez *et al*., 2019; Cristofori *et al*., 2021).

Among beneficial commensals, *Bifidobacterium* species represent prominent, gram-positive, non-motile gut inhabitants with established clinical efficacy in managing gastrointestinal and metabolic disorders (Gavsy *et al*, 2023). Specifically, *Bifidobacterium longum* (*B. longum*) has demonstrated strong potential to attenuate inflammation and oxidative stress through its active metabolic pathways and secretion of organic acids, including acetate, butyrate, propionate, and lactate (Cristofori *et al*, 2021; Chen *et al*, 2020; Gavsy *et al*, 2023). Beyond local gut barrier protection, *B. longum* modulates the gut-heart axis to regulate systemic lipid metabolism, consistently reducing total cholesterol and triglyceride concentrations (Tang *et al*, 2023).

Given the rising global prevalence of diet-induced metabolic disorders and the unique, complementary mechanisms of statins and beneficial gut microbes, integrating pharmacological and probiotic strategies represents a novel paradigm for cardiovascular risk reduction. We hypothesized that targeting the gut-heart axis via *B. longum*-linked inflammatory pathways, in combination with Atorvastatin, would offer a superior, synergistic therapeutic strategy for managing dyslipidemia, oxidative stress, and early atherogenesis. Accordingly, the present study investigates the individual and combined protective effects of Atorvastatin and *Bifidobacterium longum* against dyslipidemia, systemic redox imbalance, and tissue pathology in rats subjected to a high-cholesterol diet, providing mechanistic insights into potential translational co-therapies.

## 2 Materials required

### 2.1 Chemicals

Cholesterol (Cat# 54181) and Cholic acid (Cat# 54908) were purchased from Sisco Research Laboratories Pvt. Ltd., India (SRL). Malondialdehyde (MDA) (Cat# 63287-1G-F) and Thiobarbituric acid (TBA) (Cat# T5500-25G) were purchased from Sigma Aldrich, USA. Trichloroacetic acid (TCA) (Cat# 90544) was purchased from Sisco Research Laboratories Pvt. Ltd., India (SRL). Primary and secondary antibodies were procured from Abcam antibodies, USA; Cell Signaling technologies, USA; Thermoscientific, USA and Abclonal, China. *Bifidobacterium longum* (Strain BB536) was purchased from SuperSmart, Luxemborg. Atorvastatin (Aztor10 MG) was purchased from Sun Pharma, India

### 2.2 Animal Studies

8 weeks old Wistar male albino rats weighing around 230 ± 10 g were obtained from Central Animal House Facility, University of Madras, Taramani Campus, Chennai (Tamilnadu). The animal experiments were carried out as per the guidelines of the Committee for the Purpose of Control and Supervision of Experiments on Animals (CPCSEA), India and approved by the Institutional Animal Ethics Committee (IAEC), University of Madras, Taramani, Chennai (IAEC Approval No. 01/03/2023). Rats were acclimatized for one week under standard laboratory conditions and randomization was carried out before initiating the experiment. A High cholesterol diet (HCD) model was established by feeding a high-cholesterol diet (comprising of normal rat chow supplemented with 4% cholesterol and 1.5% cholic acid) until the end of the experimental period. Atorvastatin (20 mg/kg b.w) and *B*.*longum* (10 mg/kg b.w) were administered orally to the treatment groups for the last 20 days of the experimental period.

### 2.3 Collection of serum and tissue

At the end of the experimental period, the rats were anesthetized with ketamine (80 mg/kg body weight, i. p.) and xylazine (20 mg/kg body weight, i.p.). Blood was collected into anticoagulant-free test tubes. Blood samples were kept at room temperature for 30 min, allowed to clot and then centrifuged at 3000 rpm for 10 min to isolate the serum. The aortic and liver tissues were excised and washed in ice-cold phosphate buffered saline (PBS, pH 7.4) and stored for histological examination, the aortic tissue was fixed in 10 % neutral buffered formalin.

### 2.4 Histopathology

For histopathological evaluation, animals were anaesthetized with ketamine (22 mg□kg□1 body weight, i.p.) and aorta, liver were carefully excised. The tissues were fixed in 4% paraformaldehyde (PFA) and processed for paraffin embedding. Sections were mounted onto gelatin coated slides using PBS for subsequent Hematoxylin and eosin staining and analysis.

### 2.5 Lipid Peroxidation assessment

Serum malondialdehyde (MDA) levels was estimated according to the method described by Devasagayam and Tarachand (1987). Results were expressed as micromoles of malondialdehyde (MDA) per milligram of protein. Serum (200µL) was mixed with 10%TCA (400µL) and 1% (600µL) vortexed, and incubated at 95□°C for 30□min. After cooling, samples were centrifuged at 2000rpm for 10 min and absorbance was measured at 535nm.

### 2.6 Analysis of inflammatory markers and cell adhesion molecules by Enzyme linked immunosorbent assay

Serum concentrations of inflammatory markers (TNF-α, IL-6, IL-8, CRP) and cell adhesion molecules (VCAM-1, ICAM-1) were quantified using enzyme-linked immunosorbent assay (ELISA). Briefly, 100 μL of serum containing 25 μg of protein was coated onto 96-well plates using bicarbonate buffer and incubated overnight at 4□°C. The following day, plates were washed with phosphate-buffered saline (1X PBS) and blocked with 1% bovine serum albumin (BSA; 200 μL/well) for 2 h to minimize non-specific binding. After three washes with 1X PBS, wells were incubated overnight at 4□°C with specific primary antibodies (Table S1) (1:1000 dilution in 1% BSA; 100 μL/well). Subsequently, plates were washed four times with 1X PBS containing Tween-20 and incubated with horseradish peroxidase (HRP)-conjugated secondary antibody (Table S1) (100 μL/well) for 1 h at room temperature. After three washes with 1X PBS, 100 μL of 1 mM ABTS [2,2′-azino-bis (3-ethylbenzothiazoline-6-sulfonic acid)] substrate was added and incubated in the dark for 20 min. The reaction was terminated by adding 100 μL of 2 N sulphuric acid per well. Absorbance was measured at 415 nm using a Bio-Rad iMark™ Microplate Absorbance Reader (Bio-Rad Laboratories, Hercules, CA, USA).

## 3. RESULTS

### 3.1 HCD induced alterations in blood parameters and tissues

To investigate the therapeutic efficacy of *Bifidobacterium longum* (B), and it’s combined administration with Atorvastatin (AS), rats were subjected to a high cholesterol diet for 120 days (Figure 1A). Treatment regimens (*B. longum*, or combination therapy) were initiated on day 90 and continued through day 120 (Figure 1A).

**Figure 1.**
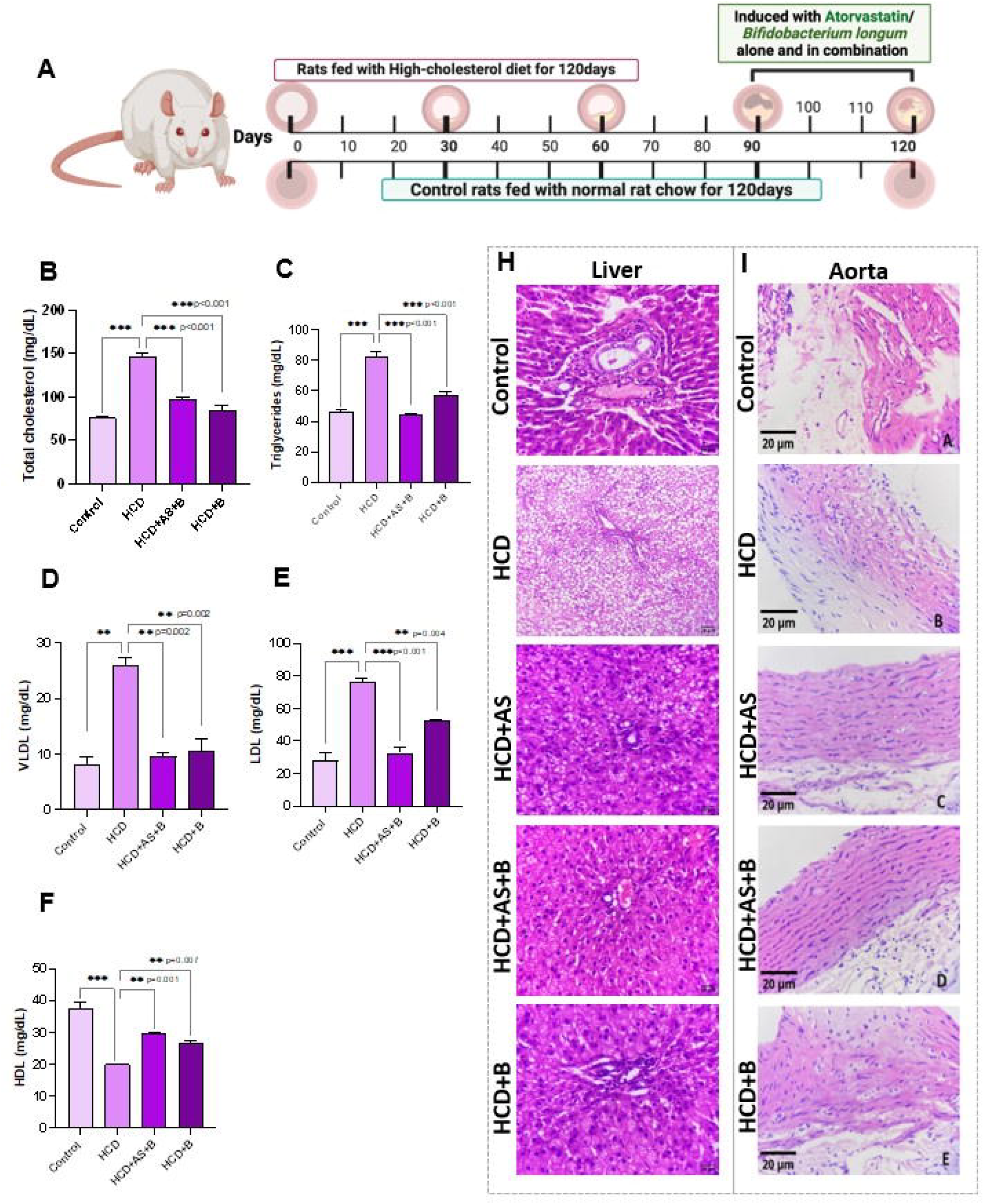
Effect of Atorvastatin and Bifidobacterium longum on lipid profile and tissue histology in HCD-fed rats.

#### 3.1.1 Effect of *B. longum* and Atorvastatin on blood glucose and Serum Lipid Profiles

HCD feeding for 120 days induced marked metabolic dysregulation in rats, characterized by significant alterations in lipid parameters when compared to the control group **(Figure 1B–F)**. Serum Total cholesterol levels were markedly elevated in the HCD group compared to controls (p<0.001). All treatment interventions *B*.*longum*, and HCD+AS + B combination showed significantly reduced total cholesterol levels compared to the HCD group (p<0.001). Notably, combination therapy (AS + B) demonstrated a superior lowering effect compared to B alone treatment (p<0.01). HCD feeding induced a robust increase in serum TG and VLDL levels (p<0.001vs. Control). Treatment with B, or combination significantly suppressed TG and VLDL accumulation (p<0.001 or p<0.01). Combination therapy (HCD + AS + B) achieved the maximum suppression, restoring TG and VLDL levels (Figure 1C, D). Serum LDL levels were dramatically increased in HCD-fed rats (p<0.001). Treatment with AS + B significantly lowered LDL levels (p<0.001), demonstrating the most pronounced reduction (Figure 1E). Conversely, HCD intake caused a substantial decline in cardioprotective HDL cholesterol (p<0.001) when compared to control). Combination treatment (HCD + AS + B) and *B. longum* monotherapy significantly boosted HDL concentrations (p<0.01) when compared to HCD rats (Figure 1F).

#### 3.1.2 Histopathological Evaluation of Liver and Aorta Tissue

To assess structural changes resulting from HCD feeding and therapeutic treatments, histological analyses of liver and aorta tissues were performed (Figure 1H, I). **Hepatic Histology:** Liver sections from the Control group exhibited normal hepatic lobular architecture with well-defined hepatocytes and sinusoids (Figure 1H, panel A). HCD feeding induced severe hepatic steatosis, characterized by extensive lipid droplet accumulation, micro-and macro-vesicular vacuolation, and loss of cellular structure (Figure 1H, panel B). Treatment with AS alone partially alleviated fat deposition (Figure 1H, panel C), whereas *B. longum* alone and combination therapy (AS + B) markedly restored liver microarchitecture, markedly reducing lipid accumulation and cellular vacuolation (Figure 1H, panels D and E). **Aortic Histology:** Cross-sections of the aorta in control rats demonstrated intact endothelial lining and smooth, organized vascular architecture (Figure 1I, panel A). HCD feeding resulted in significant vascular remodeling, endothelial disruption, and luminal thickening indicative of early atherosclerotic changes (Figure 1I, panel B). Administration of AS, B, or their combination (AS + B) effectively protected vascular integrity, maintaining normal tunica media architecture and endothelial smoothness (Figure 1I, panels C–E).

### 3.2 Modulation of Redox Dynamics and Lipid Peroxidation Markers Under High-Cholesterol Diet and Combination Therapies

To evaluate the impact of *Bifidobacterium longum*, and the combination (AS+B) on oxidative stress induced by a high-cholesterol diet (HCD), redox status and lipid peroxidation markers were evaluated (Figure 2). Schematic illustration shows the interplay between systemic oxidative stress, HDL-mediated anti-atherogenic defense, and the projected protective role of Atorvastatin and *Bifidobacterium longum* co-treatment in mitigating atherogenesis (Figure 2A).

**Figure 2.**
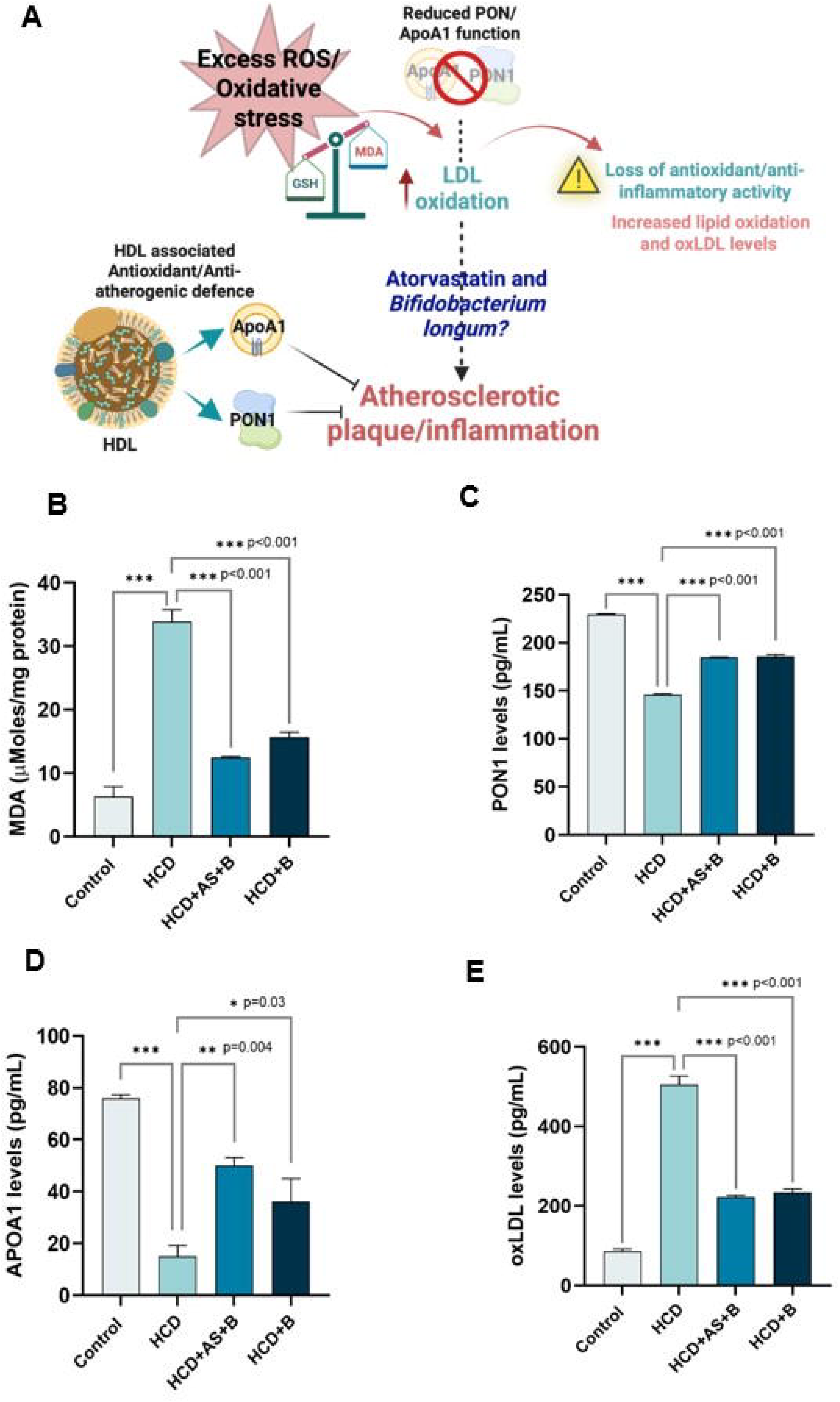
Redox Dynamics and Antioxidant Panel.

#### 3.2.1 Redox panel

Serum **malondialdehyde (MDA) Levels, a** key indicator of lipid peroxidation, **demonstrated** a robust, highly significant elevation on HCD fed rats when compared to the control group (p<0.001). Treatment with AS monotherapy significantly reduced MDA levels compared to the untreated HCD group (p<0.001). Furthermore, combination therapy (HCD + AS + B) and *B. longum* monotherapy (HCD + B) produced even greater reductions in MDA (p<0.001), restoring redox homeostasis (Figure 2B).

#### 3.2.2 Enhancement of HDL-Associated Proteins and Reduction of Oxidized LDL

The functional capacity of HDL-associated anti-atherogenic proteins (PON1 and APOA1) and systemic LDL oxidation were further analyzed in these experimental groups (Figure 2A, C–E). Serum PON1 levels were significantly depleted in HCD-fed rats compared to controls (p<0.001). Administration of *B. longum* alone, or the combination (AS + B) significantly restored PON1 levels compared to the HCD fed group (p<0.001; Figure 2C). Similarly, HCD intake caused a drastic reduction in serum APOA1 levels (p<0.001 vs. Control). Monotherapy with AS (p<0.05) and *B. longum* (p<0.05) partially recovered APOA1 expression, whereas combination therapy (AS + B) exhibited the strongest restorative effect (p<0.01) when compared to HCD fed rats (Figure 2D). Furthermore, Serum oxLDL levels were markedly elevated in the HCD group compared to controls (p<0.001). Remarkably, combination therapy (AS + B) and *B. longum* monotherapy further suppressed oxLDL levels (p<0.001 vs. HCD), substantially mitigating pro-atherogenic oxidative modifications (Figure 2E).

### 3.3 Attenuating HCD-Induced Vascular Inflammation through Atorvastatin and *Bifidobacterium longum* treatment

To elucidate the mechanisms governing vascular protection, systemic inflammatory mediators and endothelial activation markers were quantified across all experimental groups (Figure 3). Schematic representation details the cellular and molecular mechanisms through which a High-Cholesterol Diet (HCD) triggers systemic dyslipidemia, endothelial dysfunction, and vascular inflammation, ultimately driving atherogenesis **(Figure 3A)**. High Cholesterol Diet triggers the endothelial activation and pro-inflammatory cytokine secretion led to atherosclerotic progression. Excessive circulating oxidized lipoproteins upregulate transcription factor, that results in inclined levels of pro-inflammatory cytokines (CRP, TNF-α, IL-6 and IL-8) and cell adhesion molecules (ICAM-1 and VCAM-1) that facilitate leukocyte migration into the vascular endothelium. C-reactive protein (CRP) levels were significantly elevated in HCD-fed rats compared to controls (p<0.001), reflecting widespread systemic inflammation (Figure 3B). Notably, *B. longum* monotherapy (p<0.05) and combination therapy (AS + B; p<0.01) led to even more robust reductions in CRP compared to the untreated HCD group (Figure 3B). Treatment with atorvastatin significantly reduced TNF-α levels attributable to its inhibition of HMG-CoA reductase, which not only lowers cholesterol but also suppresses NF-κB mediated pro-inflammatory cytokine transcription. Treatment with AS + B (p<0.01), or *B. longum* alone (p<0.001) significantly declined TNF-α production compared to HCD (Figure 3C). IL-6 levels were significantly decreased in all the three treated groups (B, and AS + B; p<0.001) when compared to HCD fed rats, bringing cytokine levels back toward baseline (Figure 3D). Serum IL-8 concentrations were markedly reduced across all treated groups (B, and AS + B) compared to the HCD group (p<0.001; Figure 3E). Endothelial activation was evaluated via circulating levels of adhesion molecules responsible for leukocyte recruitment (Figure 3F, G). HCD feeding caused a significant upregulation of both ICAM-1 and VCAM-1 compared to control rats (p<0.001). Co-treatment with *B. longum* (AS + B) achieved the maximum reduction (p<0.001 vs. HCD), while *B. longum* monotherapy also significantly lowered ICAM-1 levels (p<0.01 vs. HCD; Figure 3F). Serum VCAM-1 levels were significantly attenuated by *B. longum* (p<0.001), and combination administration (p<0.001) relative to the HCD group, indicating effective reduction of endothelial-leukocyte adhesion cascades (Figure 3G). Altogether, these findings highlight the beneficial impact of dual pharmacological–probiotic intervention in attenuating systemic inflammation and vascular dysfunction, underscoring its potential in preventing atherosclerotic progression.

**Figure 3.**
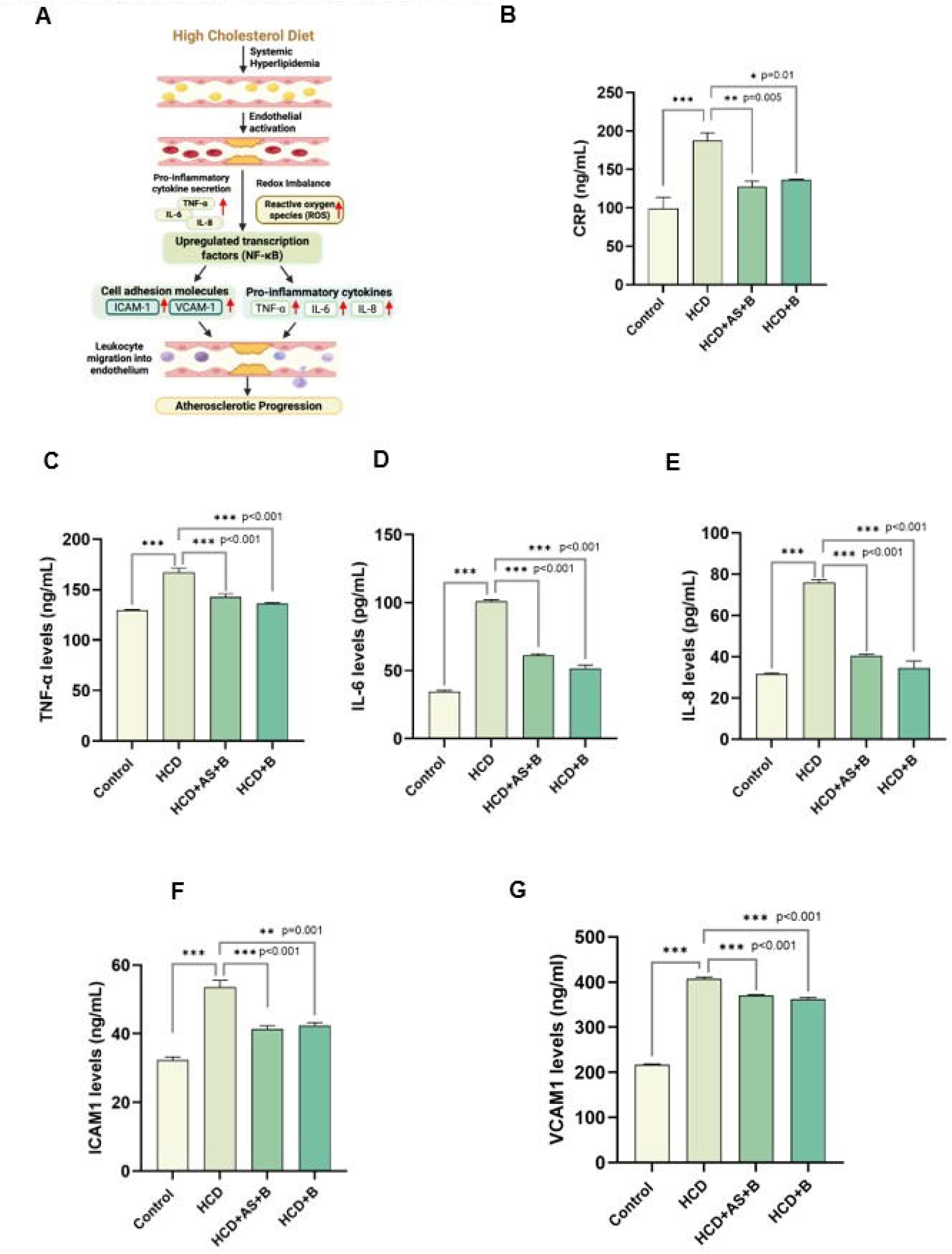
Anti-inflammatory and vascular endothelial effects of Atorvastatin and *Bifidobacterium longum* in HCD-fed rats.

## 4. Discussion

The escalating prevalence of hypercholesterolemia and its associated cardiovascular complications remains a major global public health concern. Increased LDL-C has persisted as a leading modifiable risk factor and closely linked to the development and progression of atherosclerotic cardiovascular disease (Vaduganathan *et al*., 2022). High-cholesterol diets accelerate metabolic impairment, trigger oxidative stress, and promote atherogenesis through systemic endothelial dysfunction and dyslipidemia (Jonsson and Bäckhed, 2016). While statins like Atorvastatin are the gold standard for managing hypercholesterolemia, prolonged high-dose statin therapy is frequently limited by adverse effects, including myopathy, liver enzyme elevations and development of new onset Type II Diabetes Mellitus (Ward *et al*., 2019; Mach *et al*., 2020; Stroes *et al*., 2015). Consequently, combination strategies incorporating gut-microbiota targeted therapies such as *Bifidobacterium longum* have emerged as promising adjuncts to enhance therapeutic efficacy while potentially reducing requisite statin dosages. High-cholesterol diet (HCD) feeding for 120 days established hypercholesterolemia, marked by severe dyslipidemia, glycemic elevation, systemic oxidative stress, inflammation in circulation and histological damage in hepatic and aortic tissues. Treatment with Atorvastatin, *B. longum*, or their combination effectively counteracted these diet-induced metabolic abnormalities, with combination therapy yielding superior lipid-lowering and antioxidant benefits.

### 4.1. Amelioration of Lipid Profile and Glycemic Homeostasis

High-cholesterol intake overburdens hepatic lipid transport and clearance pathways, leading to atherogenic dyslipidemia, a common finding in patients with cardiometabolic disease (Vekic *et al*, 2023). Results demonstrates that administration of Atorvastatin or *B. longum* significantly attenuated HCD-induced elevations in TC, TG, LDL, and VLDL. Atorvastatin exerts its primary therapeutic effect by inhibiting 3-hydroxy-3-methylglutaryl-coenzyme A (HMG-CoA) reductase, thereby upregulating LDL receptor expression and enhancing peripheral lipid clearance (Mach *et al*., 2020; Fan and Watanabe, 2021). Conversely, probiotics like *B. longum* modulate lipid metabolism through distinct gut-mediated pathways. Notably, the combination of Atorvastatin and *B. longum* achieved maximum suppression across TC, TG, VLDL, and LDL levels, while simultaneously promoting a substantial elevation in cardioprotective HDL levels. This synergy suggests a multi-target mechanism wherein statin-mediated inhibition of cholesterol synthesis operates in tandem with probiotic-driven cholesterol excretion and intestinal absorption blockade highlighting the broader metabolic benefits of gut microbiota modulation in diet-induced metabolic syndrome.

### 4.2. Mitigation of Oxidative Stress and Restoration of HDL Functional Capacity

Overproduction of Reactive Oxygen Species (ROS) by endothelial cells and their subsequent interaction with sub-endothelial LDL has been shown in *in-vitro* studies to be a key atherogenic iniatior (Violi and coworkers, 2024). Oxidative stress is intimately linked with hypercholesterolemia; overwhelming endogenous antioxidant defenses and causing lipid peroxidation (Loffredo and Carnevale 2024; Tsimikas and Witztum, 2024). Malondialdehyde (MDA), a major secondary product of lipid peroxidation (Ayala and coworkers, 2014), was drastically elevated in HCD-fed rats, confirming severe cellular oxidative damage. Treatment with Atorvastatin, *B. longum*, and particularly their combination, dramatically reduced MDA accumulation, underscoring strong systemic antioxidant activity. Concurrently, glutathione (GSH) levels, which spiked in the HCD group as an initial adaptive compensatory response to oxidative overload, normalized following treatment as ROS production diminished and redox homeostasis was restored.

A high influx of LDL-C, exceeding the cell’s ability efflux cholesterol promotes the formation of foam cells (Sorci-Thomas & Thomas, 2016). Beyond overall redox balance, the structural and functional integrity of HDL particles plays a critical role in anti-atherogenic defense. HDL exerts anti-inflammatory and antioxidant actions primarily through associated enzymes and structural apolipoproteins, specifically Paraoxonase 1 (PON1) and Apolipoprotein A1 (APOA1). PON1 degrades oxidized lipids in LDL and cell membranes, while APOA1 constitutes 70% of the protein content in HDL and facilitates reverse cholesterol transport (Denimal, 2023; Linton *et al*., 2023; Durrington & coworkers, 2025; Xu *et al*, 2022). In HCD-fed rats, severe depletion of PON1 and APOA1 compromised HDL functional capacity, directly facilitating the accumulation of pro-atherogenic oxidized LDL (oxLDL).

Treatment interventions most prominently the combination of Atorvastatin and *B. longum* effectively restored PON1 and APOA1 expression, leading to a profound suppression of oxLDL accumulation. By preventing LDL oxidation, this therapeutic combination directly inhibits a critical initiating step in foam cell formation, vascular inflammation, and plaque development.

High-Cholesterol Diet (HCD) induces endothelial activation and a robust pro-inflammatory cascade (). Results revealed that excessive circulating oxidized lipoproteins act as upstream stimuli that upregulate key transcription factors (such as NF-κB). This molecular shift drives the overproduction of critical systemic inflammatory mediators, including **CRP, TNF-**α, **IL-6, and IL-8**, alongside cellular adhesion molecules **ICAM-1 and VCAM-1**. The significant elevation of C-reactive protein (CRP) in HCD-fed rats (p<0.001) underscores a state of widespread, chronic systemic inflammation.

This environment promotes leukocyte recruitment and migration into the vascular endothelium, establishing the foundational cellular environment required for atherosclerotic plaque progression (Libby *et al*., 2021).

A central finding of this investigation is the efficacy of both monotherapies and dual therapy in attenuating HCD-induced vascular dysfunction. The marked reduction in TNF-α, IL-6, IL-8, and adhesion molecules by Atorvastatin alone is attributable to its established pleiotropic effects. Beyond its primary role as an HMG-CoA reductase inhibitor for lowering cholesterol levels, Atorvastatin suppresses NF-κB-mediated transcription. This downstream effect directly halts the production of pro-inflammatory cytokines and leukocyte-recruiting adhesion cascades.

Interestingly, *B. longum* monotherapy demonstrated robust anti-inflammatory capacity, achieving a statistically significant reduction in CRP and bringing IL-6 and IL-8 levels. Probiotics are increasingly recognized for their ability to modulate the gut-vascular axis. *B. longum* likely exerts these systemic benefits by reinforcing gut barrier integrity, reducing metabolic endotoxemia (LPS translocation), and producing short-chain fatty acids (SCFAs) that systemically suppress macrophage activation (). The dual pharmacological-probiotic approach yielded the most profound improvements. Most notably, the combination achieved the **maximum reduction in ICAM-1 levels** and a highly robust downregulation of CRP. This suggests a multi-targeted, synergistic therapeutic effect: while Atorvastatin works intracellularly to block hepatic lipid synthesis and endothelial NF-κB pathways, *B. longum* works via the gastrointestinal tract to minimize systemic inflammatory triggers. Together, they form a comprehensive defense against endothelial-leukocyte adhesion.

## Conclusion

In conclusion, the concurrent administration of *Bifidobacterium longum* with Atorvastatin produces a synergistic therapeutic effect against high-cholesterol diet-induced metabolic dysregulation. This combination strategy comprehensively restores serum lipid homeostasis, enhances HDL anti-atherogenic functionality (via PON1 and APOA1 upregulation), suppresses LDL oxidation, and protects hepatic and vascular tissues from lipotoxic and oxidative damage. These findings highlight probiotic co-supplementation as a compelling, low-risk strategy to enhance statin efficacy and improve clinical outcomes in dyslipidemia and cardiovascular disease management.

By targeting distinct but intersecting pathological pathways, the co-administration of Atorvastatin and *B. longum* effectively neutralizes the adhesion molecule cascades (ICAM-1 and VCAM-1) that facilitate early-stage atherogenesis. These findings provide compelling evidence that integrating gut-microbiome-targeted probiotics with standard statin therapy enhances vascular protection. This dual approach offers a promising, highly effective paradigm for mitigating chronic systemic inflammation and arresting the progression of atherosclerotic cardiovascular disease.

## Acknowledgements

The authors express their gratitude to the Department of Health Research - Multidisciplinary Research Unit (DHR-MRU) at the Dr. ALM Post Graduate Institute for Basic Medical Sciences, University of Madras, Taramani Campus, Chennai, TN, India for providing access to its state-of-the-art facilities.

