## Supplemental Table 1 for "Combined statin–probiotic treatment mitigates atherogenic dyslipidemia fueling NAFLD in high-cholesterol-diet fed rats"

**SUPPLEMENTARY DATA**

**Table S1:List of primary and secondary antibodies used for ELISA in the study**

| S. No | Antibodies | Dilution | Vendor | Identifiers |
|---|---|---|---|---|
| 1. | Mouse monoclonal anti- CRP | 1:2000 | Santa Cruz Biotechnology | Cat# sc-73864, RRID: AB_2085782 |
| 2. | Mouse monoclonal anti-IL-6 | 1:2000 | Santa Cruz Biotechnology | Cat# sc-57315, RRID: AB_2127596 |
| 3. | Mouse monoclonal anti-IL-8 | 1:1000 | Santa Cruz Biotechnology | Cat# sc-376750, RRID: AB_2891256 |
| 4. | Mouse monoclonal anti-TNF α | 1:2000 | Santa Cruz Biotechnology | Cat#sc-133192, RRID: AB_1567355 |
| 5. | Rabbit monoclonal anti-APOA1 | 1:2000 | Abcam antibodies | Cat# ab52945 RRID: AB_869970 |
| 6. | Mouse monoclonal anti-PON1 | 1:2000 | Abcam antibodies | Cat# ab24261 RRID: AB_303304 |
| 7. | Mouse monoclonal anti-OxLDL | 1:2000 | Abcam antibodies | Cat# ab17591) RRID: AB_302496 |
| 8. | Rabbit monoclonal Anti-VCAM1 | 1:2000 | Cell Signaling Technologies | Cat# 13662s RRID: AB_2798866 |
| 9. | Rabbit monoclonal Anti-ICAM1 | 1:2000 | Cell Signaling Technologies | Cat# 67836s RRID: AB_2798867 |
| 16. | Goat pAb to rabbit IgG H&L (HRP) | 1:10000 | Abcam antibodies | Cat# ab205718, RRID: AB_2819166 |
| 16. | Peroxidase AffiniPure™ Goat Anti-Rabbit IgG (H+L) | 1:5000 | Jackson Immunoresearch Laboratories Inc | Cat# 111-035-003, RRID: AB_231356 |
| 17. | Peroxidase AffiniPure™ Goat Anti-Mouse IgG (H+L) | 1:5000 | Jackson Immunoresearch Laboratories Inc | Cat# 115-035-003, RRID: AB_10015289 |
